# Translatomic profiling of the acute stress response: It’s a TRAP

**DOI:** 10.1101/2020.11.24.392464

**Authors:** Lukas von Ziegler, Johannes Bohacek, Pierre-Luc Germain

## Abstract

The impact of stress on gene expression in different cell types of the brain remains poorly characterized. Three pioneering studies have recently used translating ribosome affinity purification followed by RNA sequencing (TRAP-seq) to assess the response to stress in CA3 pyramidal neurons of the hippocampus. The results suggest that acute stress alters the translation of thousands of genes in CA3 pyramidal neurons, and that this response is strongly modulated by factors such as sex, genotype and a history of early life stress. However, our reanalysis of these datasets leads to different conclusions. We confirm that acute stress induces robust translational changes in a small set of genes. However, we found no evidence that either early life stress or sex have an effect on gene translation induced by acute stress. Our findings highlight the need for additional studies with adequate sample sizes and proper methods of analysis to assess the impact of stress across cell types in the brain.

---

Understanding the impact of stress on brain function requires a detailed knowledge of the molecular events that unfold in response to stressful experiences [1, 2]. Transcriptomic techniques have revealed some of the molecular cascades triggered by acute stressors, mostly in the mouse hippocampus, yet these studies have used bulk tissue for their analyses [3–5]. More recently, several studies have begun to address cell-type specificity of the stress response. Three pioneering studies have employed translating ribosome affinity purification (TRAP) technology [6] to characterize which genes are actively translated in CA3 pyramidal neurons of the hippocampus in response to stress [7–9]. These studies suggest that acute stress alters the translation of thousands of genes in CA3 pyramidal neurons, and that this response is strongly modulated by factors such as sex, genotype and a history of early life stress. However, a careful reanalysis of these data - which comprise two datasets deposited at Gene Expression Omnibus (GSE100579 and GSE131972) – leads to different conclusions. Here, we present a re-analysis and an updated interpretation of these important data.

The first study [7] reports that in CA3 pyramidal neurons, thousands of genes respond differently to acute stress (forced swim test – FST) in males and females (Figure 1A). However, our re-analysis indicates that, most likely due to the low number of replicates (6 independent samples for 4 groups) and high technical variability (Figure 1C), this does not replicate in samples from the second dataset (Figure 1B). A follow-up study [8] reports that early life stress (ELS) changes the translational response of a large number of genes to an acute stress challenge later in life. To demonstrate this, the sets of genes that were (significantly) changed in response to acute stress in the ELS group were subtracted from those changed in the non-ELS control group. Visual inspection of the genes reveals highly variable effects (Figure 1D). The appropriate method to test whether ELS impacts the translational response to an acute stressor is to use interaction terms in generalized linear models. We implemented this approach with different packages, including the one used by the authors in the original publication, as well as a (more lenient) method based on staged false discovery rate calculations [10]. These analyses fail to yield a single gene showing significant effect of ELS on the stress response. A detailed description and full documentation of our reanalysis is publicly available: https://github.com/ETHZ-INS/stressTRAPreanalysis.

**Figure 1:**
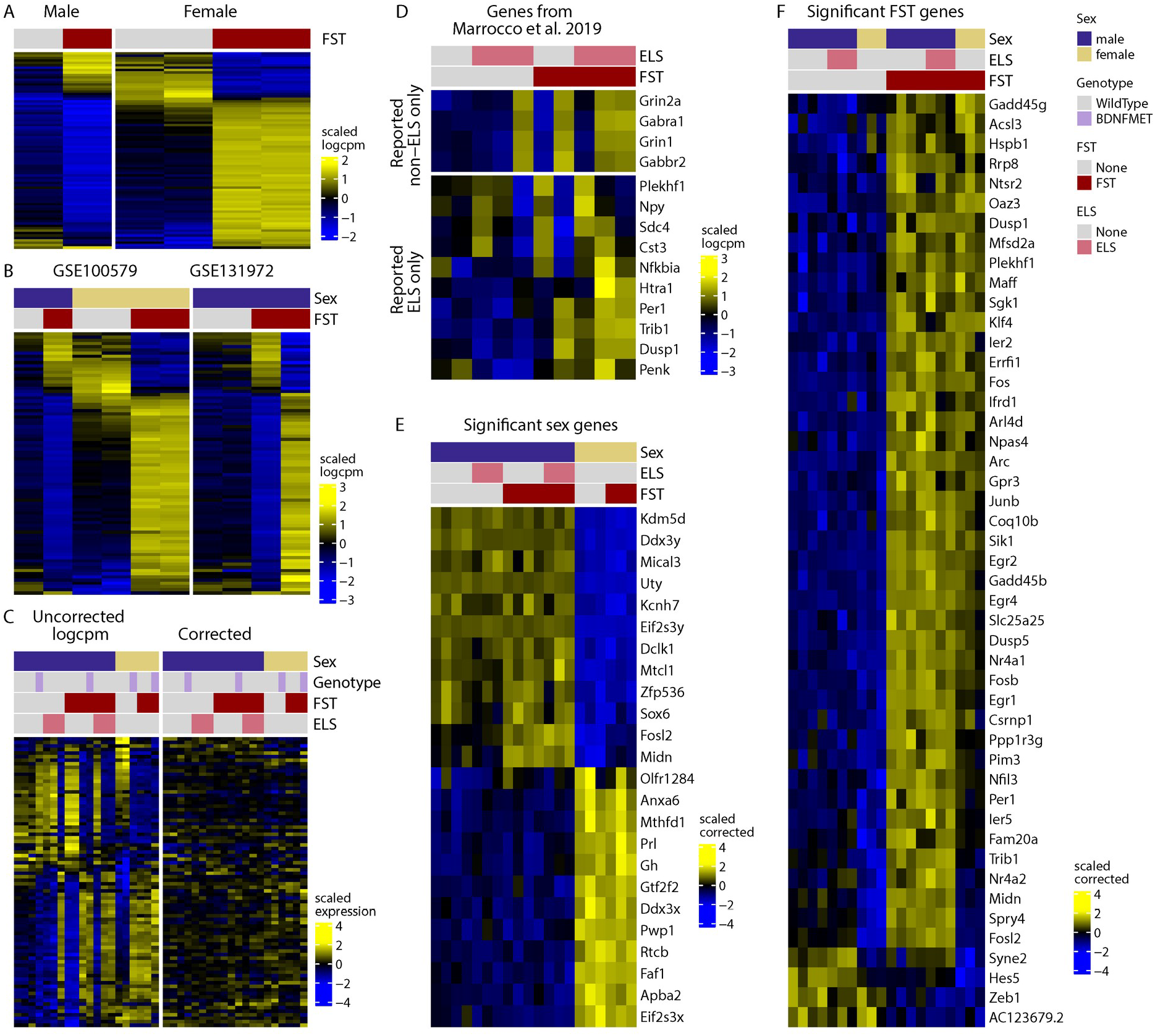
Reanalysis of environmentally induced changes in the translatome of CA3 pyramidal neurons. A) genes that show sex-dependent responses to stress as reported by Marrocco et al. (2017, data from GSE100579 wild-type mice). B) Including non-ELS data from GSE131972 shows that these genes are highly variable in the male-FST group, suggesting that the sex-dependent effects observed in the first dataset were spurious effects due to random variation. C) Including all data highlights the strong co-expression of these genes in a manner that is not explained by experimental groups (left panel). Removing technical variability eliminates the apparent sex-specific signal (right panel), suggesting that these differences were an artefact. D) Genes reported by Marrocco et. al (2019) to be responsive to FST only in non-ELS (top) or ELS (bottom) animals. E) Genes that are differentially translated in females and males across all data in our global reanalysis. F) Genes that are translationally altered following FST across all data according to our global reanalysis. The heatmap colors represent variance-scaled log-normalized expression values (i.e. row z-scores).

Reasoning that all datasets could, together, shed more light on group differences, we performed a global reanalysis of the data. We identified robust baseline translation differences in 24 genes between males and females in CA3 neurons (Figure 1E). We also found that an acute swim stress challenge induces robust translational changes in 47 genes (Figure 1F), compared to >2000 genes originally reported by Gray et al [9]. However, we found no evidence that either sex or ELS impact the translational response to acute stress, nor that ELS persistently alters gene translation. Finally, in a third study [9] hundreds of genes were reported to respond to acute stress only in the stress-hypersensitive BDNF-Val66Met mouse model, but not wild-type mice. Our reanalysis reveals only a subtle impact of genotype on a few genes (for detailed results, see https://github.com/ETHZ-INS/stressTRAPreanalysis).

In summary, our reanalysis reveals that acute stress induces translational changes in a small set of genes in CA3 pyramidal neurons (Figure 1F), many of which are well-known, stress-sensitive immediate early genes also detected in bulk RNA-sequencing after stress [5]. We find no evidence that either ELS or sex have an effect on gene translation induced by acute stress in CA3 pyramidal neurons. Importantly, this does not prove the absence of such effects. Rather, it demonstrates that even when aggregating all data these studies are still underpowered, requiring yet more replicates to draw solid conclusions. As new multi-omics techniques rapidly emerge, it is important to remember that the high initial costs associated with sophisticated new techniques, such as TRAP-seq, ATAC-seq or single-cell sequencing, often leads to strongly underpowered studies. Pooling samples is no alternative to proper biological replication, and in the absence of sufficient data, results of such studies must be interpreted with caution.

## Acknowledgements

The lab of JB is funded by the ETH Zurich, ETH Project Grant ETH-20 19-1, SNSF Grant 310030_172889/1, the Botnar Foundation and the Swiss 3R Competence Center. The position of PLG is co-funded by Prof. Isabelle Mansuy (Brain Research Institute, University Zürich and Institute of Neuroscience, ETH Zurich), Profs. Gerhard Schratt and Johannes Bohacek (Institute of Neuroscience, ETH Zurich) and by Prof. Mark Robinson (Institute of Molecular Life Sciences, University of Zurich).

